# RNase H1 and H2 are differentially regulated to eliminate RNA-DNA hybrids

**DOI:** 10.1101/593152

**Authors:** Arianna Lockhart, Vanessa Borges Pires, Fabio Bento, Vanessa Kellner, Sarah Luke-Glaser, Brian Luke

## Abstract

RNA-DNA hybrids are tightly regulated to ensure genome integrity. The RNase H enzymes, RNase H1 and H2, contribute to chromosomal stability through the removal of RNA-DNA hybrids. Loss of RNase H2 function is implicated in human diseases of the nervous system and cancer. To better understand RNA-DNA hybrid dynamics, we have focused on elucidating the regulation of the RNase H enzymes themselves. Using yeast as a model system, we demonstrate that RNase H1 and H2 are controlled in different manners. RNase H2 is regulated in a strict cell cycle dependent manner, both in terms of its R-loop removal, and ribonucleotide excision repair functions. RNase H1, however, can function independent of cell cycle stage to remove R-loops, but appears to become activated in response to high R-loop loads. These results provide us with a more complete understanding of how and when RNA-DNA hybrids are acted upon by the RNase H enzymes.

## INTRODUCTION

The RNase H enzymes (RNase H1 and RNase H2) remove RNA-DNA hybrids through the endonucleolytic cleavage of an RNA moiety that is engaged in the hybrid molecule (Cerritelli and Crouch, 2009). R-loops are a form of RNA-DNA hybrid that arise when an RNA transcript base-pairs to its DNA template, resulting in displacement of the non-coding DNA strand (Santos-Pereira and Aguilera, 2015). Defective R-loop removal results in a loss of genome integrity, likely due to DNA replication conflicts (Garcia-Muse and Aguilera, 2016). RNase H1, coded by the *RNH1* gene in yeast, is able to degrade RNA engaged in R-loops, and its overexpression is frequently used as a tool to promote their removal (Huertas and Aguilera, 2003; Wahba et al., 2011). RNase H1 consists of an N-terminal hybrid binding domain (HBD) and a C-terminal endonuclease motif (Cerritelli and Crouch, 2009) the latter of which must interact with the 2’-OH group of four consecutive ribose molecules for optimal activity (Nowotny et al., 2007). RNase H1 is likely recruited to hybrids via its direct interaction with replication protein A (RPA), a single stranded binding protein that can coat the displaced DNA strand of an R-loop (Nguyen et al., 2017). *Rnaseh1*^*-/-*^ mice die during embryonic development due to incomplete mitochondrial replication (Cerritelli et al., 2003), a function not conserved in budding yeast due to the lack of a mitochondrial targeting sequence (MTS) on the yeast protein (Arudchandran et al., 2000). Surprisingly, the single deletion of *RNH1* in yeast has only subtle growth defects. A study by Zimmer and Koshland demonstrated that loss of heterozygosity (LOH) occurrs to the same extent in *rnh1* cells as in wild type cells (Zimmer and Koshland, 2016). Using a chromatin immunoprecipitation (ChIP) based approach, the Zimmer and Koshland study demonstrated that although Rnh1 is able to associate with R-loops across the genome, it is only active at a small subset of those loci; particularly at strong R-loop forming loci (Zimmer and Koshland, 2016). The authors suggest that Rnh1 may be constitutively negatively regulated and only undergo activation in response to R-loop mediated stress, i.e. when R-loops become strongly stabilized. Overexpression would potentially overcome the constraints of endogenous Rnh1 regulation, which would explain the ability of increased Rnh1 levels to effectively process R-loops genome-wide. Alternatively, or additionally, Rnh1 activity may be regulated in a cell cycle-dependent manner, although an initial study has demonstrated that mRNA levels of *RNH1* do not fluctuate throughout the cell cycle (Arudchandran et al., 2000). The nature of Rnh1 regulation remains to be determined.

As in humans, yeast RNase H2 is composed of three subunits, Rnh201 (catalytic), Rnh202 and Rnh203. Mutations in all three human subunits (RNASEH2A, RNASEH2B and RNASEH2C, respectively) have been demonstrated to contribute to the neuro-inflammatory disease, AGS (Aicardi-Goutières syndrome) (Crow et al., 2006). Recently, *RNASEH2B* deletion was observed to frequently occur in CLL (<u>c</u>hronic lymphocytic leukemia) and CRPC (<u>c</u>astration <u>r</u>esistant <u>p</u>rostate <u>c</u>ancer) (Zimmermann et al., 2018). RNase H2, like RNase H1, is able to degrade the RNA in R-loops, however in addition, RNase H2 can perform <u>r</u>ibonucleotide <u>e</u>xcision <u>r</u>epair (RER), whereby rNMPs (ribonucleoside monophosphates) are excised from otherwise duplex DNA and the remaining nick gets subsequently repaired (Williams et al., 2016). RER is required when ribonucleotides are mistakenly incorporated into chromosomes during DNA replication by DNA polymerases. Misincorporated ribonucleotides are considered one of the most frequently occurring DNA lesions and have been estimated to occur 10,000 times /S phase/cell in budding yeast (Nick McElhinny et al., 2010a). When RER is defective, in the absence of RNase H2 function, topoisomerase I (Top1) can aberrantly process rNMPs (Williams et al., 2013), which leads to deletions and formation of double strand breaks (DSBs) (Huang et al., 2017; Kim et al., 2011). The aberrant processing of rNMPs by Top1 also leads to genome instability in human cells when RNase H2 is mutated (Zimmermann et al., 2018).

Unlike impaired RNase H1 function, the loss of RNase H2 results in a stronger genome instability phenotype in yeast (Conover et al., 2015; O’Connell et al., 2015; Zimmer and Koshland, 2016), consistent with the notion that RNase H2 accounts for the majority of RNase H activity in a cell (Arudchandran et al., 2000). It has been difficult however, to determine which enzymatic activity of RNase H2 is responsible for promoting genome integrity. The generation of a separation of function mutant allele of the catalytic subunit of yeast RNase H2 (*RNH201-P45D-Y219A*) that is <u>r</u>ibonucleotide <u>e</u>xcision repair <u>d</u>efective, hence *RNH201-RED,* has shed some light on this matter (Chon et al., 2013). Using this allele, it has been suggested that the majority of genomic defects in yeast associated with the loss of RNase H2 activity are due to faulty R-loop processing and not defective RER (Conover et al., 2015; O’Connell et al., 2015; Zimmer and Koshland, 2016). Nonetheless, expression of the *RED* allele did not completely suppress LOH phenotypes in the Zimmer and Koshland study, indicating that unprocessed rNMPs may also contribute to chromosomal aberrations. ChIP experiments demonstrated that RNase H1 was detected at R-loops, whereas RNase H2 was found to only weakly associate with R-loops, and its distribution remains unclear (Conover et al., 2015; O’Connell et al., 2015; Zimmer and Koshland, 2016). At telomeres, where R-loops are present, Rnh201 has been shown to accumulate only very late in S phase (Graf et al., 2017). Consistently, the mRNA expression of *RNH201* shows peaks in S and G2 phases of the cell cycle, suggesting a potential cell cycle regulated activity (Arudchandran et al., 2000).

In this study, we have attempted to better understand how the RNase H enzymes are regulated, with respect to how and when they remove RNA-DNA hybrids from the genome. By creating cell cycle restricted alleles of the RNase H enzymes, we were able to demonstrate that RNase H2 is able to support all of its functions, i.e. R-loop removal and RER, in the G2 phase of the cell cycle, but not during the S phase. In contrast, RNase H1 was able to act at R-loops irrespective of cell cycle stage, but appeared to require an R-loop induced stress to trigger its activity, as has been speculated by Zimmer and Koshland (Conover et al., 2015; O’Connell et al., 2015; Zimmer and Koshland, 2016). The cell cycle regulated activity of RNase H2 is reflected in terms of its stronger association with chromatin/telomeres in the G2 phase. Together, the results provide us with a better understanding with respect to how RNA-DNA hybrid removal is accomplished by the RNase H enzymes, which may have implications for diseases associated with RNA-DNA hybrid misregulation.

## RESULTS

### The RNase H enzymes can be expressed in a cell cycle dependent manner

We set out to address whether the RNase H enzymes are regulated, in terms of their functional capacity, during the cell cycle. As a starting point, we employed endogenous tandem affinity purification (TAP)-tagged versions of RNase H1 (Rnh1-TAP) as well as all three subunits of RNase H2 (Rnh201-TAP, Rnh202-TAP and Rnh203-TAP) expressed under their native promoter (Figure 1A, top). We have previously demonstrated that TAP tagged versions of RNase H1 and H2 are fully functional (Graf et al., 2017). To follow protein expression levels, yeast cells were synchronized in the G1 phase with α-factor and released into the cell cycle at 25°C, with samples for western blotting and DNA content analysis being collected every 15 minutes. We observed that levels of Rnh1-TAP (Figure 1B, top), Rnh202-TAP (Figure 1C, top) and Rnh203-TAP (Figure S1A) were stable throughout all phases of the cell cycle. The expression of the catalytic subunit of RNase H2 (Rnh201-TAP) was slightly enhanced following the bulk of DNA replication as compared to other cell cycle stages (Figure S1B). DNA staining coupled to flow cytometry was used to ensure that cell synchronization and release protocols had properly functioned for RNase H1 and all subunits of RNase H2 (Figure S1C and S1D). In addition, protein membranes were probed for the Sic1 and Clb2 proteins, which are expressed in G1 and G2/M, respectively (Figures 1B, 1C and S1A, S1B). In order to interrogate the cell cycle contribution of the RNases H, we created alleles of *RNH1* and *RNH202* in which expression of the proteins was restricted to either the S or G2 phase of the cell cycle. For the S phase specific expression of RNase H1 and H2, we introduced the promoter of the S phase induced cyclin, *CLB6*, along with an in-frame fusion of the Clb6 degron (Hombauer et al., 2011) to the 5’ end of the *RNH1* (*S-RNH1-TAP*) and *RNH202* (*S-RNH202-TAP*) genes (Figure 1A, middle panel). Yeast cells were synchronized in G1 and released as described above. Western Blotting of samples taken throughout the cell cycle revealed that the S-tagged versions were now no longer expressed evenly throughout the cell cycle, but rather peaked in S-phase, between Sic1 degradation and Clb2 expression, as expected (Figures 1B, 1C middle panels). G2/M expressed alleles of the same genes (*G2-RNH1-TAP* and *G2-RNH202-TAP*) were created using a similar strategy, but with the promoter and degron sequences of Clb2 (Hombauer et al., 2011) (Figure 1A bottom). Accordingly, the G2-tagging resulted in a restricted expression to the G2 and M phases of the cell cycle (Figure 1B, 1C bottom panels), concomitant with the G2 and mitotic marker Clb2. Expression of *RNH1* and *RNH202* in either S-phase or the G2/M phase did not affect unperturbed cell cycle progression as assayed by flow cytometry-based analysis of DNA content (Supplementary Figures S1C and S1D). Importantly, although RNase H2 is a trimeric protein, cell cycle specific expression of one subunit (in our case Rnh202) should be sufficient to confine the enzyme’s activity to a particular cell cycle stage, as the holoenzyme requires all three subunits for its activity (Cerritelli and Crouch, 2009). In summary, we have created cell cycle restricted alleles of the RNase H enzymes, which will assist in understanding how they contribute to the removal of RNA-DNA hybrids.

**Figure 1.**
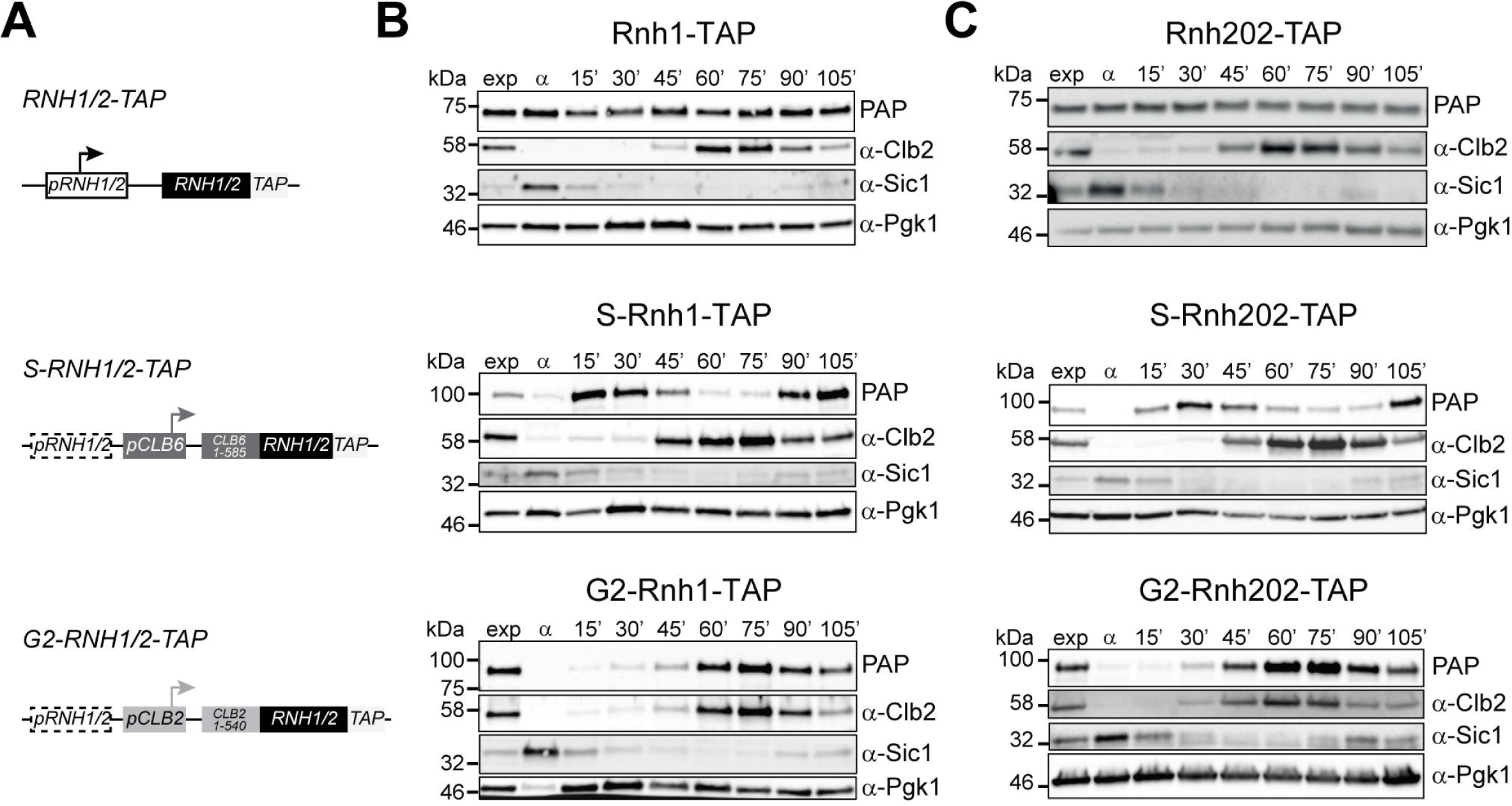
Expression of RNases H in a cell cycle dependent manner. (A) Schematic of the RNases H genomic loci (top) and of the modified S phase (middle) and G2 (bottom) expressed RNase H alleles. (B and C) Western blot analysis showing Rnh1, S-Rnh1 and G2-Rnh1 (B) and Rnh202, S-Rnh202 and G2-Rnh202 (C) protein levels in the cell cycle. Exponentially growing cells were arrested in G1 with α-factor and released synchronously in the cell cycle at 25°C. Protein samples were collected at 15 min intervals. See also Figure S1.

### R-loop removal by RNase H2 is critical post replicatively

*rnh1 rnh201* double mutants are sensitive to the DNA damaging agent methyl methanesulfonate (MMS) (Lazzaro et al., 2012). By spotting serial dilutions of yeast onto solid media with or without MMS, we were able to confirm this result (Figure 2A, third row from bottom). The expression of a wild-type copy of *RNH201* rescues the MMS sensitivity, as expected (Figure 2A). Importantly, the *RNH201-P45D-Y219A* allele, a mono-<u>r</u>ibonucleotide <u>e</u>xcision repair <u>d</u>efective allele (hereafter referred to as ‘RED’) (Chon et al., 2013), could also rescue the MMS sensitivity of *rnh1 rnh201* double mutants, albeit not quite to the same extent as the wild type copy. This result indicates that the MMS sensitivity of RNase H mutant cells is largely linked to the faulty removal of R-loops and not misincorporated ribonucleotides. Accordingly, we observed that the addition of MMS to both wild type and *rnh1 rnh201* mutants increased the amount of R-loops, as detected by dot blot analysis with the S9.6 antibody, which recognized RNA-DNA hybrids (Figures 2B and 2C).

**Figure 2.**
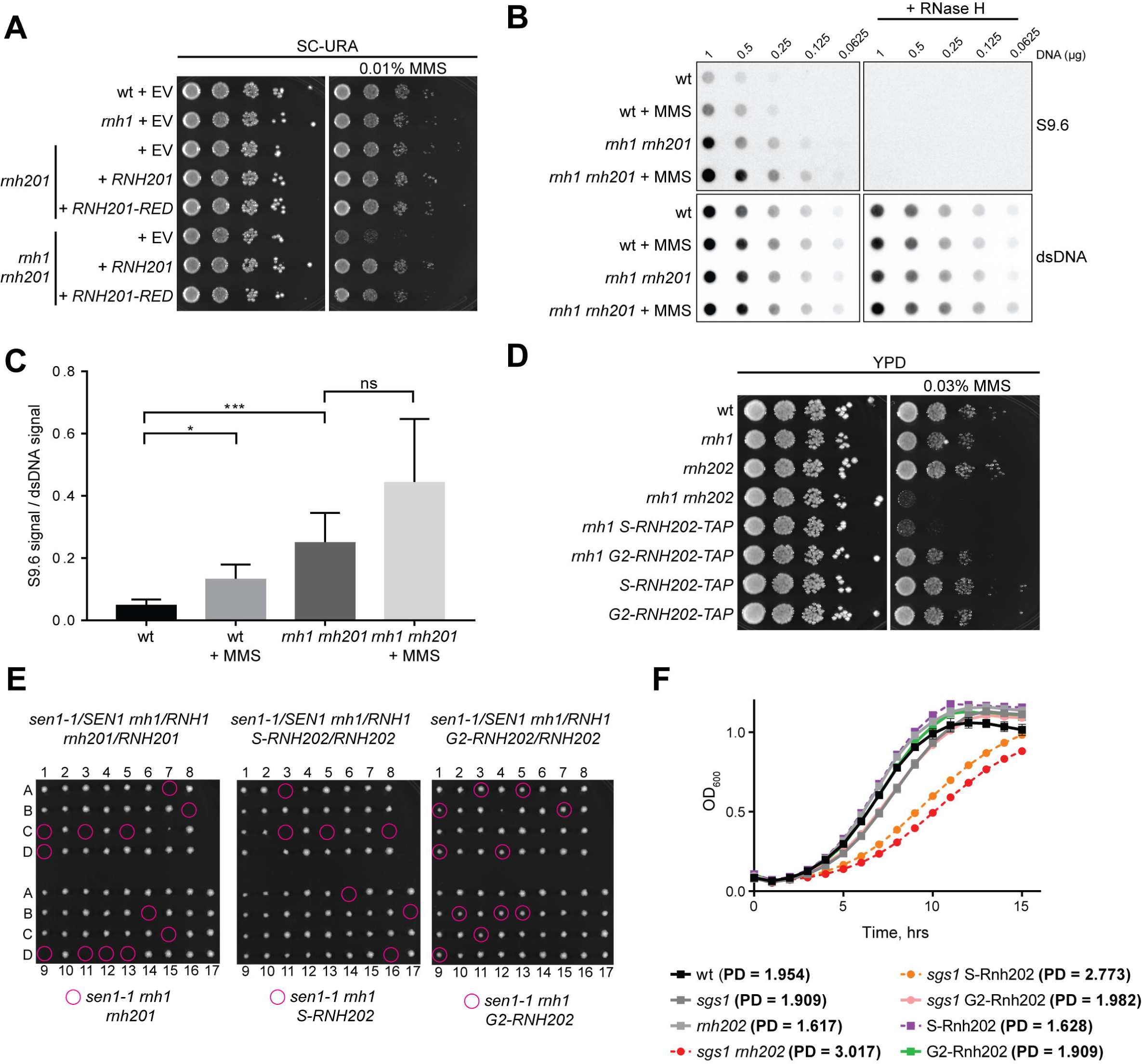
R-loop removal by RNase H2 is critical post-replicatively. (A) A tenfold serial dilution of the indicated strains transformed with the indicated plasmids was spotted onto SC-URA plates with or without 0.01% MMS. Images were taken after 2 days growth at 30°C. EV = empty vector. (B) Dot blot analysis of R-loop levels in the indicated strains, either exponentially growing or treated with 0.02% MMS for 90 minutes. All samples were treated with RNase III and RNase T1, and as a control they were additionally treated with Rnase H. Membranes were incubated with S9.6 or α-dsDNA antibodies. (C) Quantification of (B), indicating S9.6 signal normalized to dsDNA signal. Mean values + SEM of 3 biological replicates are shown for each strain. p values were obtained from two-tailed paired Student’s t tests (*p<0.05; ***p<0.001; ns, not significant). (D) A tenfold serial dilution of the indicated strains was spotted onto rich medium with or without 0.03% MMS. Images were taken after 2 days growth at 30°C. (E) Tetrad analysis of the meiotic products from the indicated heterozygous diploid strains. Diploids were micromanipulated onto rich medium and plates were grown for 4 days at 25°C. Genotypes were confirmed by replica plating and scoring for markers linked to the respective mutations (see yeast strains for markers). Magenta circles correspond to triple mutant spores of the indicated genotype. (F) Kinetic growth analysis of the indicated strains. Cells were inoculated at 0.05 OD_600_ in SC complete and grown for 15 hours at 30°C in 96 well microtiter plates. OD_600_ measurements were performed every hour. Mean values ± SEM of 3 biological replicates are shown for each strain. In brackets, the calculated population doubling (PD) time in hours of each strain is indicated. See also Figure S2.

Similar to *rnh1 rnh201* mutants, we also observed that *rnh1 rnh202* double mutants were sensitive to MMS (Figure 2D). Strikingly, the expression of *S-RNH202-TAP* failed to rescue the MMS sensitivity in a *rnh1* mutant background, whereas the *G2-RNH202-TAP* promoted growth similar to *rnh1* single mutants (Figure 2D). The combined results suggest that RNase H2 has a critical role in R-loop removal in a post-replicative manner. To corroborate that the R-loop removal activity of RNase H2 is critical in the G2 phase of the cell cycle, we employed a hypomorphic/temperature-sensitive allele of the RNA-DNA hybrid helicase Sen1 (*sen1-1*) (Mischo et al., 2011), in the context of the RNase H2 cell cycle alleles. It has recently been described that cell death ensues in *rnh1 rnh201 sen1-1* mutants due to the accumulation of toxic R-loops (Costantino and Koshland, 2018). Indeed, we attempted to generate triple mutants by sporulating a heterozygous diploid strain and did not recover any viable triple mutants (Figure 2E, left). We were also unable to recover viable *sen1-1 rnh1 S-RNH202-TAP* mutants, however the *sen1-1 rnh1 G2-RNH202-TAP mutants* were viable, and spores were indistinguishable from wild type (Figure 2E, middle and right), supporting the conclusion that RNase H2 acts in G2 to remove R-loops.

Loss of the yeast Bloom and Werner helicase homolog, *SGS1,* displays a synthetic growth defect specifically with the loss of RNase H2, due to the faulty removal of R-loops and not due to faulty RER (Chon et al., 2013). In an *sgs1* background, the *S*-*RNH202-TAP* expressing cells grew with similar kinetics as *RNH202* deletion mutants, whereas the *G2-RNH202-TAP* expressing cells divided with similar kinetics as a wild type *RNH202* allele in the same background (Figure 2F). Identical relationships between *sgs1* and the cell cycle alleles of RNase H2 can be observed in terms of MMS sensitivity as well (Figure S2). Taken all together, these results indicate the important role of RNase H2 activity in R-loop removal in G2 and further suggest that RNase H2 R-loop removal activity is inefficient during the S phase.

### RNase H2 performs RER post-replicatively

Misincorporated rNMPs accumulate when RNase H2 activity is impaired, leading to faulty RER. This is exacerbated when a steric gate mutant allele of the catalytic subunit of DNA polymerase ε(*pol2M644G*) is expressed, which incorporates 10x more rNMPs (Nick McElhinny et al., 2010b) and renders the cells sensitive to hydroxyurea (HU). *rnh201 pol2M644G* cells exposed to HU can be rescued through the expression of wild type RNase H2, but not the RED allele, as previously shown, (Lafuente-Barquero et al., 2017) (Figure 3A), demonstrating the specificity of the HU sensitivity for faulty RER in this genetic context. The G2-restricted expression of *RNH202* is able to fully rescue the HU sensitivity in the presence of the *pol2M644G* allele, whereas S phase restriction of *RNH202* expression resulted in slightly greater HU sensitivity than the *rnh202 pol2M644G* double mutants (Figure 3B).

**Figure 3.**
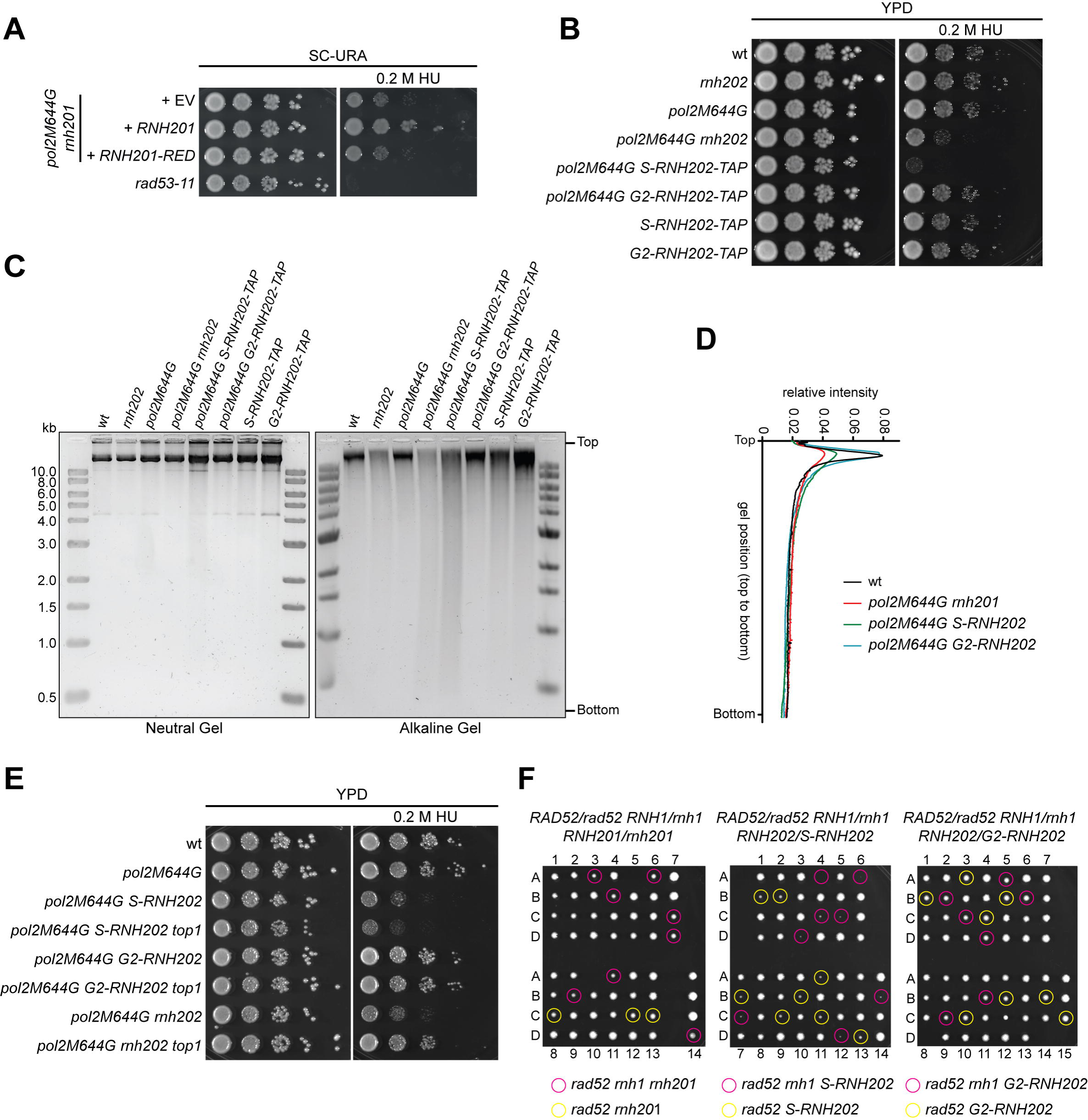
RNase H2 performs RER post-replicatively. (A) A tenfold serial dilution of the yeast strains transformed with the indicated plasmids was spotted onto SC-URA plates with or without 0.2 M HU. Images were taken after 2 (no drug) or 3 (0.2 M HU) days growth at 30°C. EV = empty vector. (B) A tenfold serial dilution of the indicated strains was spotted onto rich medium with or without 0.2 M HU. Images were taken after 2 (no drug) or 3 (0.2 M HU) days growth at 30°C. (C) Detection of alkaline-sensitive sites in genomic DNA. Genomic DNA of the indicated strains was loaded on a neutral gel after KCl treatment (left) or an alkaline gel after KOH treatment (right). Gels were stained with SyBr Gold. (D) Densitometry analysis of the data from the alkaline gel in (C). (E) A tenfold serial dilution of the indicated strains was spotted onto rich medium with or without 0.2 M HU. Images were taken after 2 (no drug) or 3 (0.2 M HU) days growth at 30°C. (F) Tetrad analysis of the meiotic products from the indicated heterozygous diploid strains. Diploids were micromanipulated onto rich medium and plates were grown for 3 days at 30°C. Magenta circles correspond to triple mutant spores (with *RNH1* deletion) and yellow circles correspond to the indicated double mutant spores that harbor either the *rnh201* deletion, the *S-RNH202* allele, or the *G2-RNH202* allele.

To confirm that the G2 expressed allele is actually performing RER we indirectly assayed the amount of rNMP insertions through the analysis of genomic DNA on both neutral and alkaline gels. DNA with a high rNMP content is more readily cleaved in alkaline conditions causing a “smearing” down the gel and hence loss of intensity of the major (intact) genomic DNA band, as can be observed in the *rnh202 pol2M644G* mutants (Nick McElhinny et al., 2010a) (Figure 3C, 3D). Whereas expression of the *G2-RNH202* allele was able to efficiently remove genomic rNMPs, the *S-RNH202* was not and resembled the profile of the *rnh202 pol2M644G* cells (Figures 3C, 3D). A large extent of the genome instability generated from faulty RER, in the absence of RNase H2 activity, is due to the error prone processing of rNMPs by Top1, which leads to non-ligatable breaks, slippage mutations and the formation of double strand breaks (DSBs) (Huang et al., 2017; Potenski et al., 2014; Williams et al., 2013). As expected, the HU sensitivity of *rnh202 pol2M644G* mutants was rescued by the further deletion of the *TOP1* gene (Figure 3E, compare bottom two rows). It was surprising to see that the deletion of *TOP1* was no longer able to rescue the HU sensitivity of the *pol2M644G* allele when *S-RNH202* was expressed (Figure 3E, compare rows 3 and 4). This suggests that the S phase restricted expression of RNase H2 may be inducing DNA damage in a Top1-independent manner. Indeed, the expression of *S-RNH202,* was particularly toxic in the absence of *RAD52,* both in the absence (magenta) or presence (yellow) of RNase H1, suggesting that homology directed repair (HDR) becomes essential when RNase H2 activity is restricted to S phase (Figure 3F).

### RNase H2 associates with chromatin following bulk DNA replication

The above results indicated that the G2 expressed form of RNase H2 is sufficient to execute both its RER and R-loop removal functions, whereby the S restricted allele is not. Since the RNase H2 subunits are expressed during all phases of the cell cycle, this suggest some form of regulation with respect to its activity. We fractionated chromatin associated- (C) and soluble (S) -proteins in cells progressing synchronously through the cell cycle to address when Rnh201-TAP was recruited to chromatin. We were unable to detect a chromatin association in either G1 or early S phase, where Rnh201-TAP remained soluble, however following the bulk of DNA replication, at the 60 minute time point following G1 release, we were able to detect Rnh201-TAP in the chromatin fraction (Figure 4A, 4B). Fractionation efficiency was controlled by re-probing the western blots for PGK1 and H3, which appear in the soluble and chromatin bound fractions, respectively. Therefore, the ability of RNase H2 to fulfill all of its RNA-DNA hybrid removal functions in G2, and not in S, is likely related to the temporal regulation of its chromatin association. This is consistent with the previously published telomere localization of Rnh201-TAP by ChIP (Graf et al., 2017).

**Figure 4.**
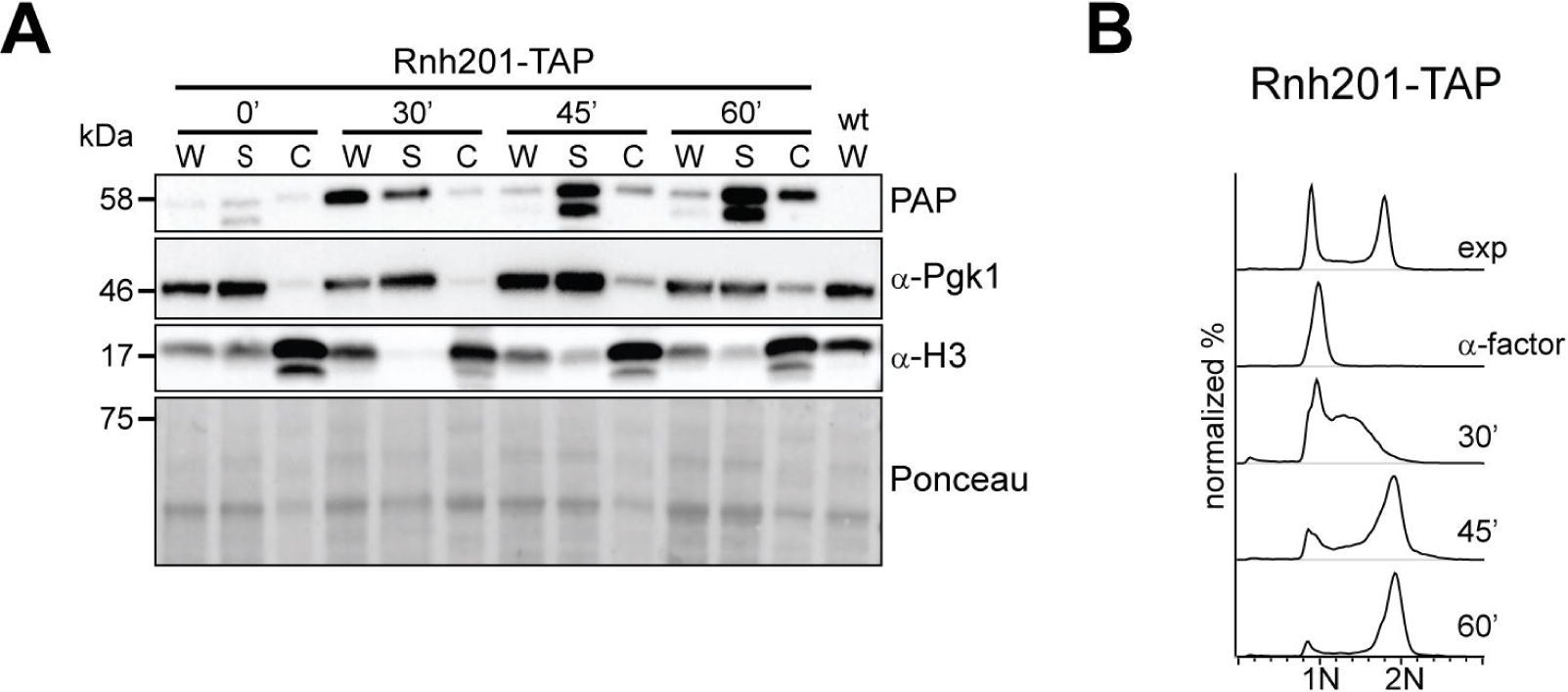
RNase H2 associates with chromatin following bulk DNA replication. (A) Chromatin association of Rnh201 in the cell cycle. Exponentially growing cells were arrested in G1 with α-factor and released in the cell cycle at 25°C. Protein samples were collected at the indicated timepoints and whole cell extract (W), soluble (S) and chromatin (C) fractions were analyzed by western blotting. Chromatin fractions were loaded 4x relative to (S) and (W). (B) Flow cytometry analysis of DNA content of samples shown in A.

### RNase H1 removes RNA-DNA hybrids when stress arises

RNase H1 activity does not contribute significantly to RER, especially when RNase H2 activity is present (Cerritelli and Crouch, 2009). Consistently, we do not observe HU sensitivity in either *rnh1 pol2M644G* cells or in the S- or G2-expressed alleles of *RNH1* in the *polM644G* background, and the overexpression of *RNH1* is not able to rescue the HU sensitivity of the *rnh202 pol2M644G* mutant (Figure S3A, S3B). On the other hand, RNase H1 is efficient in the removal of R-loops (Cerritelli and Crouch, 2009). Upon analysis of the meiotic products of heterozygous diploids, we observed that, unlike RNase H2, both the S and G2 alleles of RNase H1 were able to rescue the synthetic lethality of *rnh1 rnh201 sen1-1* mutants (Figure 5A). Similarly, both the *S-* and *G2-RNH1* alleles could grow in combination with the *RNH201* deletion on MMS plates (Figure 5B). RNase H1 was weakly, but equally, chromatin bound in all cell cycle stages (Figures 5C, 5D), consistent with a lack of obvious cell cycle regulation. Finally, the restricted expression of Rnh1 in either the S or G2 phases did not lead to lethality in a *rad52* background (Figure S3C), in contrast to the S phase restriction of RNase H2. Taken together these results suggest that RNase H1 is not cell cycle regulated, nor is the cell cycle regulation required for optimal growth.

**Figure 5.**
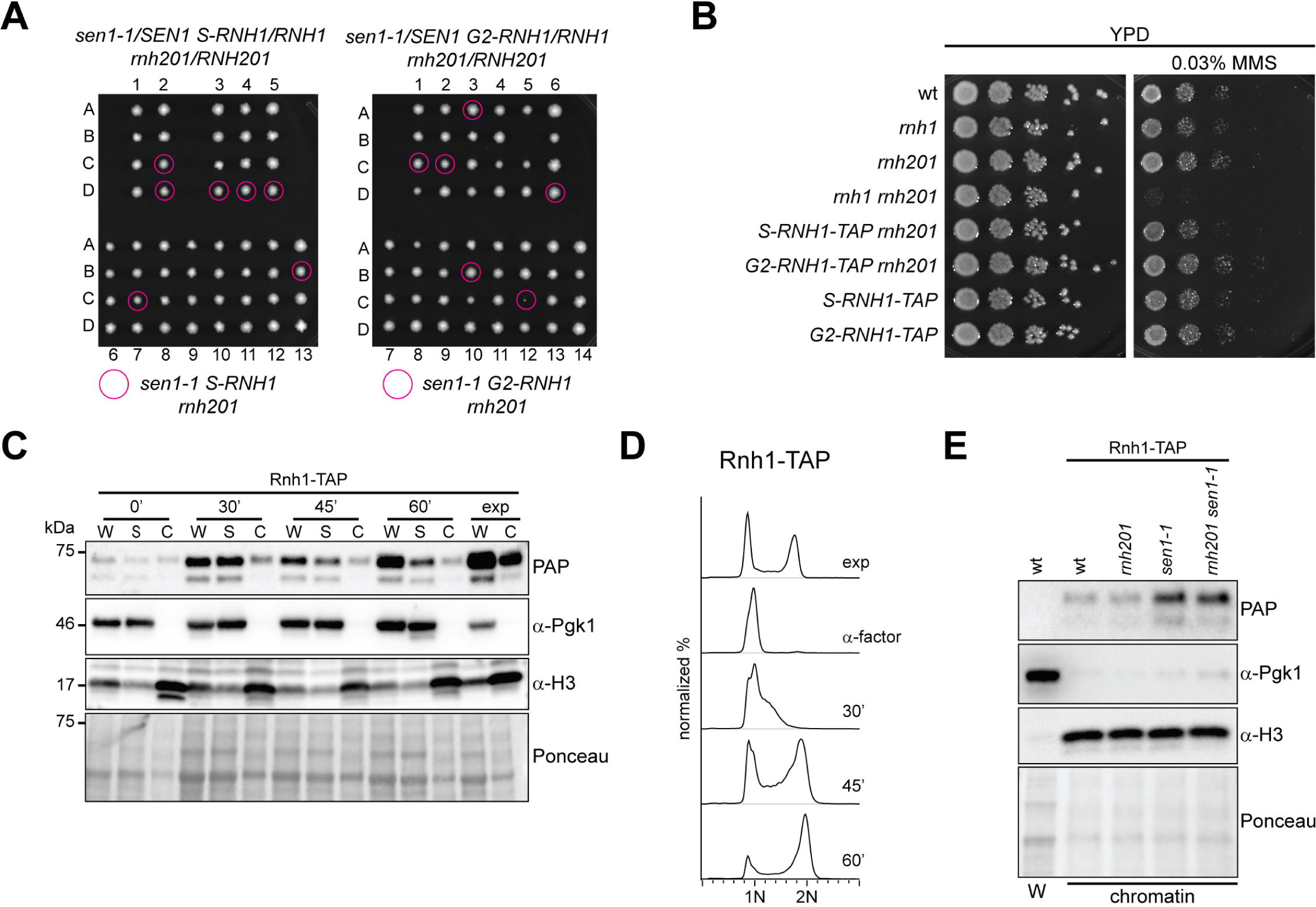
RNase H1 removes RNA-DNA hybrids when stress arises. (A) Tetrad analysis of the meiotic products from the indicated heterozygous diploid strains. Diploids were micromanipulated onto rich medium and plates were grown for 3 days at 25°C. Magenta circles correspond to triple mutant spores. (B) A tenfold serial dilution of the indicated strains was spotted onto rich medium with or without 0.03% MMS. Images were taken after 2 days growth at 30°C. (C) Chromatin association of Rnh1 in the cell cycle. Exponentially growing cells were arrested in G1 with α-factor and released in the cell cycle at 25°C. Chromatin binding assays were carried out as described in Figure 4. Chromatin fractions were loaded 4x relative to (S) and (W). (D) Flow cytometry analysis of DNA content of samples shown in C. (E) Chromatin association of Rnh1 in *sen1-1* mutants. Cells of the indicated genotypes were grown exponentially at 25°C and then shifted at 30°C for 90 minutes. Protein samples were collected and chromatin (C) fractions were analyzed by western blotting. A whole cell extract (W) of wild type was loaded as a control. See also Figure S3.

As Zimmer and Koshland suggested that Rnh1 become activated in response to stress (Zimmer and Koshland, 2016), we evaluated the chromatin localization of Rnh1 in the context of a *sen1-1* mutant background, where R-loops drastically accumulate (Costantino and Koshland, 2018). Indeed, at the semi-permissive temperature of 30°C we detected increased levels of chromatin associated Rnh1, only when the *sen1-1* mutation was present (Figure 5E and S3D). Therefore, irrespective of cell cycle stage, Rnh1 responds to increased R-loop levels.

## DISCUSSION

The Zimmer and Koshland study speculated that the RNase H2 and H1 enzymes are regulated in terms of cell cycle and in response to R-loop associated cellular stresses, respectively (Zimmer and Koshland, 2016). This hypothesis was based on their observations that although RNase H2 provided the majority of the RNase H activity in the cell, it was only weakly associated to chromatin as assayed by ChIP. They suggested that perhaps it was difficult to ChIP RNase H2 to chromatin because it may only be associated in a certain cell cycle time point. RNase H1, on the other hand, was found associated to multiple R-loops along the chromatin, but it was not actively processing the majority of them. Since the overexpression of RNase H1 is able to degrade many R-loops, it suggested that a repressor may need to be inhibited in order to unleash RNase H1 activity and that this repressor is “titrated” out upon overexpression (Zimmer and Koshland, 2016). To test these ideas in greater detail, our study has employed cell cycle restricted alleles of RNase H1 and RNase H2 and confirmed that indeed RNase H2 is tightly cell cycle regulated. Moreover, we were able to demonstrate that RNase H1 is likely reacting to R-loop mediated stress, and this appears to occur in a cell cycle independent fashion (Figure 6).

**Figure 6.**
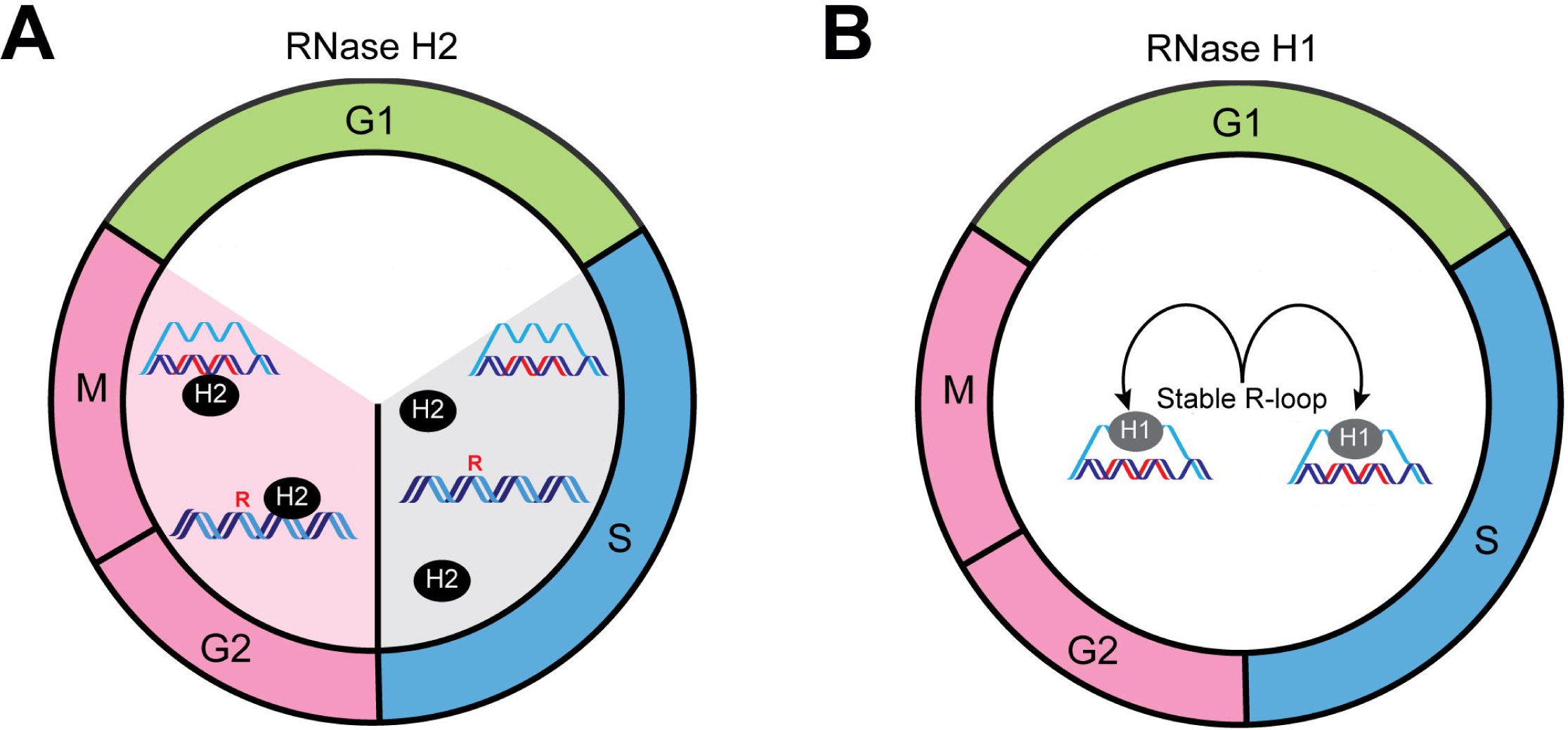

RNase H2 is able to remove RNA-DNA hybrids in R-loops as well as perform RER (Cerritelli and Crouch, 2009). MMS induces DNA damage and replication stress, both of which can lead to the stabilization and generation of R-loops (Hamperl et al., 2017; Svikovic et al., 2018; Teloni et al., 2018) (Figure 2), which likely contribute to the MMS sensitivity of *rnh1 rnh201* cells. R-loop stabilization in *sen1-1* mutants is also responsible for lethality in *rnh1 rnh201* double mutants (Costantino and Koshland, 2018). In both settings (*sen1-1* and MMS exposure), it is only the expression of RNase H2 in the G2 phase of the cell cycle that is able to support viability, whereas the S phase expression behaves similar to a deletion. This is in line with previous observations that we have made at telomeres using a ChIP assay, where RNase H2 becomes recruited to telomeres only very late in the S phase, where it presumably degrades TERRA RNA-DNA hybrids (Graf et al., 2017). Such a regulation is likely also happening on a genome-wide scale as we report here that RNase H2 only associates with chromatin once the bulk of DNA has been replicated (Figure 4). In this respect, the cell cycle restricted alleles and the chromatin/telomere association are consistent in terms of their behavior.

The RER function associated with RNase H2 is also restricted to a post-replicative time frame. Here also, the timing of rNMP removal from the genome (Figure 3) corresponds well with the chromatin association of RNase H2. It will be interesting to understand the mechanistic details with regard to how RNase H2 is regulated in the cell cycle with respect to its chromatin localization. Although the Rnh202 subunit of RNase H2 has a PCNA interacting motif (PIP box) (Chon et al., 2013) and can associate with PCNA in Co-IP experiments, we found that the removal of the PIP box was able to complement the MMS sensitivity of the *rnh202 rnh1* double mutants just as well as the wild type version (data now shown). At telomeres, we have found that the telomere associated factor, Rif2, interacts with RNase H2 and is responsible for its telomeric localization to remove TERRA R-loops (Graf et al., 2017). This however cannot account for the association of RNase H2 to other genomic loci as the *rif2 rnh1* double mutant is not sensitive to MMS like the *rnh201 rnh1* double mutant is (Graf et al., 2017). Both the Rnh202 and Rnh203 mutants have been found to harbor phosphorylated residues in phospho-proteomics approaches. It is possible that these phosphorylation events help to direct RNase H2 to the chromatin in a temporal manner (Swaney et al., 2013). Finally, the fact that RNase H2 functions primarily in a post-replicative manner to remove R-loops and excise rNMPs strongly suggests that its activity and localization may be coupled to post-replicative repair (PRR). In this respect it is curious that *TOP1* deletion is not able to rescue the S-RNase H2 allele when ribonucleotides accumulate (Figure 3). Although we do not fully understand this result at this stage, one explanation could be that exclusive expression of RNase H2 in the S phase nicks the double helix at rNMPs, and that these are converted into DSBs upon encounter with the replicating helicases. The exclusive expression of RNase H2 in S phase would then not allow the potential PRR function of RNase H2, which needs to occur in G2. Despite the fact that we do not detect RNase H2 on the chromatin in the S phase, there may be enough residual protein/activity left to make sufficient nicks for toxicity. The potential repair function of RNase H2 clearly warrants further investigation.

RNase H1 can interact with RPA, which may account for its constitutive localization to R-loops via the RPA coated displaced single stranded DNA (Nguyen et al., 2017; Petzold et al., 2015). We have not been able to detect RNase H1 at telomeres in telomerase positive cells (Graf et al., 2017), but it may be present in telomerase negative cells when telomeric R-loops accumulate and become stabilized. In any case, both the S- and G2-alleles of RNase H1 were able to support growth in *sen1-1* cells (Figure 5), suggesting that when R-loops accumulate, RNase H1 can act irrespective of cell cycle stage. This scenario was mirrored in the presence of MMS, which can induce DNA damage and replication stress, both of which can allow R-loops to accumulate and become stabilized (Hamperl et al., 2017; Teloni et al., 2018). Indeed, both the S- and G2-expressed alleles of RNase H1 could support the growth of *rnh201* cells in the presence of MMS. Together, these results reveal that that irrespective of cell cycle state, RNase H1 is able to function in scenarios where R-loops accumulate to otherwise toxic levels. Our data agrees with the interpretation from the Zimmer and Koshland study, that there is likely a switch that has to occur on RNase H1 in order to activate it when R-loops become hyper stable. This could potentially be a post-translational modification or the removal of an inhibitor protein.

We have drawn a simple model which summarizes our findings (Figure 6). In the case of RNase H2, the chromatin association is cell cycle regulated and only occurs in a post-replicative manner. Therefore, the RNA-DNA hybrids that occur in the S phase are likely only weakly processed by RNase H2. In the case of RER this restriction is probably important to minimize single strand nicking during ongoing DNA replication, which could lead to the generation of DSBs. RNase H1 on the other hand is not cell cycle regulated and appears to become activated when needed, i.e. in the case of persistent R-loops. These persistent R-loops can occur during replication stress, faulty transcription termination, or in the absence of RNase H2 function. With this in mind, our results provide insights as to why two RNase H enzymes co-exist. In theory RNase H2 should be sufficient to perform all functions required, as it can remove R-loops and perform RER. However, the activation of RER during S phase could be dangerous as DSBs may occur (see above). This may be why only RNase H1 responds to R-loop stress in the S phase, i.e. to remove the R-loop without nicking the DNA in the S phase. RNase H enzymes are critical regulators of RNA-DNA hybrids. Understanding the detailed regulation of these enzymes will contribute greatly to the general comprehension of genome maintenance with respect to RNA-DNA hybrid regulation and misregulation.

## ACKNOWLEDGMENTS

We thank: the Luke lab members for support and discussions, the media lab, Flow cytometry and protein production core facilities of the IMB and the Ulrich Lab for sharing reagents. BL’s lab was supported by the Heisenberg Program of the DFG - LU 1709/2-1. V.B.P. is funded through the GABBA PhD Program and Fundação para a Ciência e Tecnologia, Portugal (Scholarship PD/BD/127999/2016).

## AUTHOR CONTRIBUTIONS

Conceptualization, B.L., A.L., V.K.; Methodology, A.L., V.B.P., F.B.; Investigation, A.L., V.B.P., F.B., S.L.G., V.K.; Writing – Original Draft, B.L., S.L.G., A.L.; Funding Acquisition, B.L., V.B.P.; Supervision, B.L. and A.L.

## STAR METHODS

### Contact for resources and resource sharing

Further information and requests for resources and reagents should be directed to and will be fulfilled by the Lead Contact, Brian Luke.

### Experimental model and subject details

*Saccharomyces cerevisiae* strains used in this paper are derivatives of the standard strain S288C and are listed in Table S1.

Strains were grown under standard conditions in YPD (Yeast Peptone Dextrose) or SC without amino acids (Synthetic Complete) at 30°C if not indicated otherwise. Further specifications are mentioned within the Method Details section.

### Method details

#### Construction of cell cycle regulated RNase H enzymes

The *S*- and *G2*-*RNH1-TAP* alleles were created by amplifying the “*S* cassette” (containing the *CLB6* promoter, the first 585 bp of *CLB6* and the *NAT* resistance cassette) from the plasmid pBL491 with the oligonucleotides pair oAL47 + oAL48, or the “*G2* cassette” (containing the *CLB2* promoter, the first 540 bp of *CLB2* including the L26A mutation, and the *NAT* resistance cassette) from the plasmid pBL492 with the oligonucleotide pair oAL47 + oAL49. Correct integration was verified by PCR and sequencing with the oligonucleotide pairs oAL53 + oAL54 and oAL53 + oBL29. The *S-* and *G2-RNH202-TAP* alleles were created by amplifying the *S* and G2 cassettes from the plasmids pBL491 and pBL492, respectively, with the oligonucleotide pairs oAL61 + oAL62 and oAL61 + oAL63, respectively. Correct integration was verified by PCR and sequencing with the oligonucleotide pairs yAL64 + yAL65 and yAL64 + oBL29.

### α-factor arrest and release

Exponentially growing mating type a cultures were treated with 4 µg/ml of α-factor (Zymo research) and incubated further for 2 h 15 min at 30°C. Cells were then spun down and washed three times with pre-warmed water (25°C), and finally resuspended in fresh pre-warmed YPD medium (25°C) and further grown at 25°C in a water bath. Protein and FACS samples were collected at the indicated timepoints.

### SDS-PAGE and Western Blot

2 OD_600_ units were resuspended in 150 µl solution 1 (1.85 M NaOH, 1.09 M 2-mercaptoethanol) and incubated for 10 min on ice. 150 µl solution 2 (50% TCA) were added before further incubation for 10 min on ice. After centrifugation, the pellet was resuspended with 1 ml acetone, and centrifuged again. The pellet was resuspended in 100 µl urea buffer (120 mM Tris-HCl pH 6.8, 5% glycerol, 8 M urea, 143 mM 2-mercaptoethanol, 8% SDS, bromophenol blue indicator) and incubated for 5 min at 75°C followed by a rapid centrifugation step. Samples were loaded on Precast gels (Bio-Rad). The following antibodies were used at the following dilutions: PAP (Sigma-Aldrich) 1:1,000; α-Pgk1 (Invitrogen) at 1:200,000; α-Clb2 (Santa Cruz) at 1:1,000; α-Sic1 (Santa Cruz) at 1:200; α-H3 (Abcam) 1:1,000. Goat anti-rabbit -HRP conjugate and Goat anti-mouse -HRP conjugate (Bio-Rad) were used as secondary antibodies 1:3,000. Proteins were detected using the Super Signal West Pico Chemiluminescent Substrate (Thermo Scientific) or the Super Signal West Femto Chemiluminescent Substrate (Thermo Scientific) on Bio-Rad ChemiDoc™ Touch Imaging System.

### Yeast Spotting Assay

Yeast cells were incubated overnight at the appropriate temperature in YPD or SC medium. Cells were diluted to 0.5 OD_600_ and spotted in ten-fold dilutions onto YPD plates, SC plates, or plates containing the indicated amount of MMS (Sigma-Aldrich) or HU (Sigma-Aldrich). Plates were then incubated at the indicated temperatures and time and imaged using the Bio-Rad ChemiDoc™ Touch Imaging System.

### DNA content flow cytometry

0.18 OD_600_ units were washed once with water and then fixed in 70% ethanol overnight at 4°C. The fixed cells were washed once in water and then resuspended in 500 µl of 50 mM Tris-Cl pH 7.5. Afterwards 10 µl of RNase A (10 mg/ml stock RNase A; Thermo Scientific) were added to each sample and RNA was degraded for 3 h at 37°C. 25 µl proteinase K (20 mg/ml stock, Qiagen) were added and the samples were incubated for 1 to 2 h at 50°C. The cells were then resuspended in 500 µl of 50 mM Tris-Cl pH 7.5 and sonicated manually for 10 s using a Branson Sonifier® 450. 500 µl of 50 mM Tris-Cl pH 7.5 supplemented with 1 µM Sytox Green (Thermo Scientific) were added before the DNA content was measured with BD FACSVerse™.

### Dot blot for R-loop levels

Genomic DNA from either exponentially growing yeast cultures or cultures treated with 0.02% MMS for 90 min at 30°C was extracted with Gentra Puregene Yeast/Bact. Kit B (Qiagen), omitting the final RNase A treatment step. 4.8 µg DNA of each sample were treated with 5 µl RNase III (Invitrogen) and 1 µl of RNase T1 (Thermo Scientific) for 2 h at 37°C, followed by inactivation for 20 min at 65°C. As a control, 4.8 µg DNA of each sample were treated with 2 µl RNase H (NEB) (in addition to RNase III and RNase T1) for 2 h at 37°C, followed by inactivation for 20 min at 65°C. Each treated sample was splitted in two and transferred into two multiwell plates, in a final volume of 240 µl in 2x SSC, and serially diluted 1:2 in 2x SSC. From each multiwell plate, 100 µl of each well was then transferred onto a Nylon positively charged membrane (Roche) using Bio-Dot™ Apparatus (Biorad). DNA was crosslinked to the membrane with UV light (auto crosslink, Stratalinker) and after blocking the membranes in 5% milk in 1x PBST, the membranes were incubated overnight either with S9.6 (1:6,000 in 3% BSA in 1x PBST) or α-dsDNA (1:1,000 in 3% BSA in 1x PBST). Goat anti-mouse HRP conjugate (Bio-Rad) was used as secondary antibody 1:3,000. R-loops and dsDNA were detected using Super Signal West Pico Chemiluminescent Substrate (Thermo Scientific) on Bio-Rad ChemiDoc™ Touch Imaging System. Dot-blots signal was quantified in ImageJ.

### Growth kinetic analysis

Yeast cells were diluted to 0.05 OD_600_ in 100 µl of SC complete and transferred to a 96-well plate (Falcon^®^). Hourly OD_600_ measurements were performed at 30°C for 15 h by using a Tecan Spark^®^ microplate reader.

### Analysis of alkaline-sensitive sites in genomic DNA

10-20 µg of genomic DNA were treated either with 0.3 M KOH or 0.3 M KCl in a final volume of 40 µl at 55°C for 2 h in the dark. 6x alkaline loading buffer (300 mM KOH, 6 mM EDTA, 18% Ficoll 400, 0.15% bromocrestol green, 0.25% xylene cyanol) was added to KOH treated samples, which were then loaded onto a 1% alkaline agarose gel (50 mM NaOH, 1 mM EDTA) in alkaline running buffer (50 mM NaOH, 1 mM EDTA). 6x neutral loading buffer (30% glycerol in TE, 0.25% bromophenol blue, 0.25% xylene cyanol) was added to KCl treated samples, which were then loaded onto a 1% agarose gel in TBE. Both gels were run at 65 V for 5 min and then at 1 V/cm for 18 h. The alkaline gel was neutralized twice by soaking in neutralization solution (1 M Tris HCl pH 8.0, 1.5 M NaCl) for 45 min, and both gels were stained with 1x SYBR Gold (Invitrogen) in water (alkaline gel) or TBE (neutral gel) for 2 h in the dark, and imaged using the Bio-Rad ChemiDoc™ Touch Imaging System. Alkaline-sensitive sites in genomic DNA were quantified in Image J. Briefly, signal intensity was measured along the indicated lanes of the alkaline treated gel. Absolute values were then normalized by the total signal intensity per lane. Data are plotted as % signal intensity relative to the position on the gel (top to bottom).

### Chromatin binding assay

Cell pellets were washed with cold Spheroblasting Buffer (1 M sorbitol, 50 mM potassium phosphate buffer pH 7.4, 1 mM DTT) and 12 OD_600_ units were resuspended in a final volume of 1 ml in Spheroblasting Buffer. 1 µl zymolyase T100 (Brand) and 1 µl DTT were added to each sample, followed by incubation for 40 min at 30°C. Spheroblasts were collected by centrifugation and were resuspended in 300 µl Extraction Buffer (50 mM Tris-HCl, 100 mM KCl, 2.5 mM MgCl_2_, protease inhibitor cocktail (Roche), phosSTOP (Roche)). Each sample was then split in whole cell extract (50 µl, corresponding to 2 OD_600_), soluble (50 µl, corresponding to 2 OD_600_) and chromatin (200 µl, corresponding to 8 OD_600_) fractions. To each sample 0.25% final concentration of Triton X-100 was added, followed by incubation for 5 min on ice. Preparation of whole cell extract fraction: 1 µl Benzonase (Sigma) was added to each sample followed by incubation for 15 min on ice. Preparation of soluble fraction: samples were centrifuged and the supernatant was transferred to a fresh tube. Preparation of chromatin fraction: 1 ml of cold 30% sucrose solution was added to each sample, followed by centrifugation. The pellet was then resuspended in 200 µl Extraction Buffer and 5 µl 10% Triton X-100. This step was repeated again and the pellet was finally resuspended in 50 µl Extraction Buffer and 0.25% final concentration of Triton X-100. 1 µl Benzonase was then added to the samples, followed by incubation for 15 min on ice. All fractions were finally resuspended in 20 µl Urea buffer (see SDS-PAGE and Western Blot section) and samples were boiled for 5 min at 95°C. Samples were then loaded onto 4-15% precast gel (Bio-Rad).

### Key resources table

#### Supplemental items

Supplemental Figures

Table S1. Yeast strains used in this study. Related to Star Methods.

Table S2. Plasmids used in this study. Related to Star Methods.

