## Supplemental Figures for "RNase H1 and H2 are differentially regulated to eliminate RNA-DNA hybrids"

### Supplemental items

#### Supplementary Figures

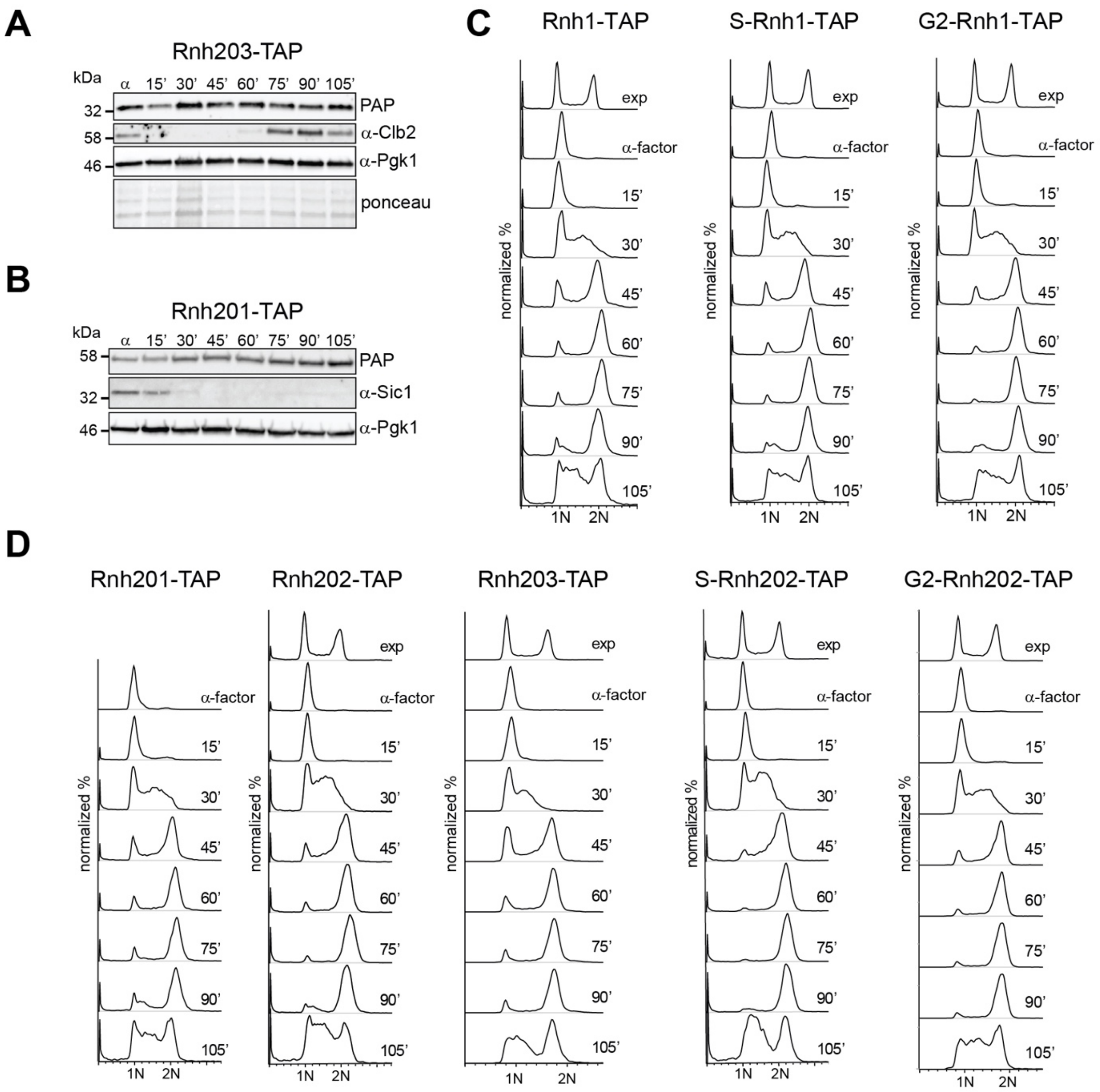


**Figure S1. Cell cycle expression of RNase H enzymes. Related to Figure 1.**

(A and B) Western blot analysis showing Rnh203 (A) and Rnh201 (B) protein levels in the cell cycle. Exponentially growing cells were arrested in G1 with α-factor and released in the cell cycle at 25˚C. Protein samples were collected at 15 min intervals.

(C) Flow cytometry analysis of DNA content of samples shown in Figure 1B.

(D) Flow cytometry analysis of DNA content of samples shown in Figure 1C and in Figure S1A and B.

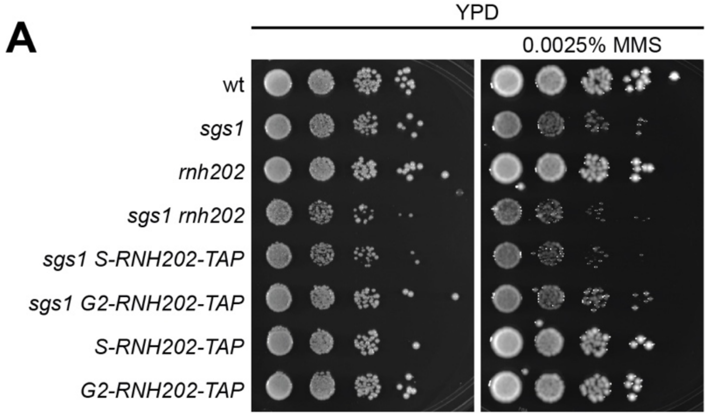


**Figure S2. RNase H2 R-loop removal activity occurs in G2 also in a *sgs1* background. Related to Figure 2.**

(A) A tenfold serial dilution of the indicated strains was spotted onto rich medium with or without 0.0025% MMS. Images were taken after 2 days growth at 30˚C.

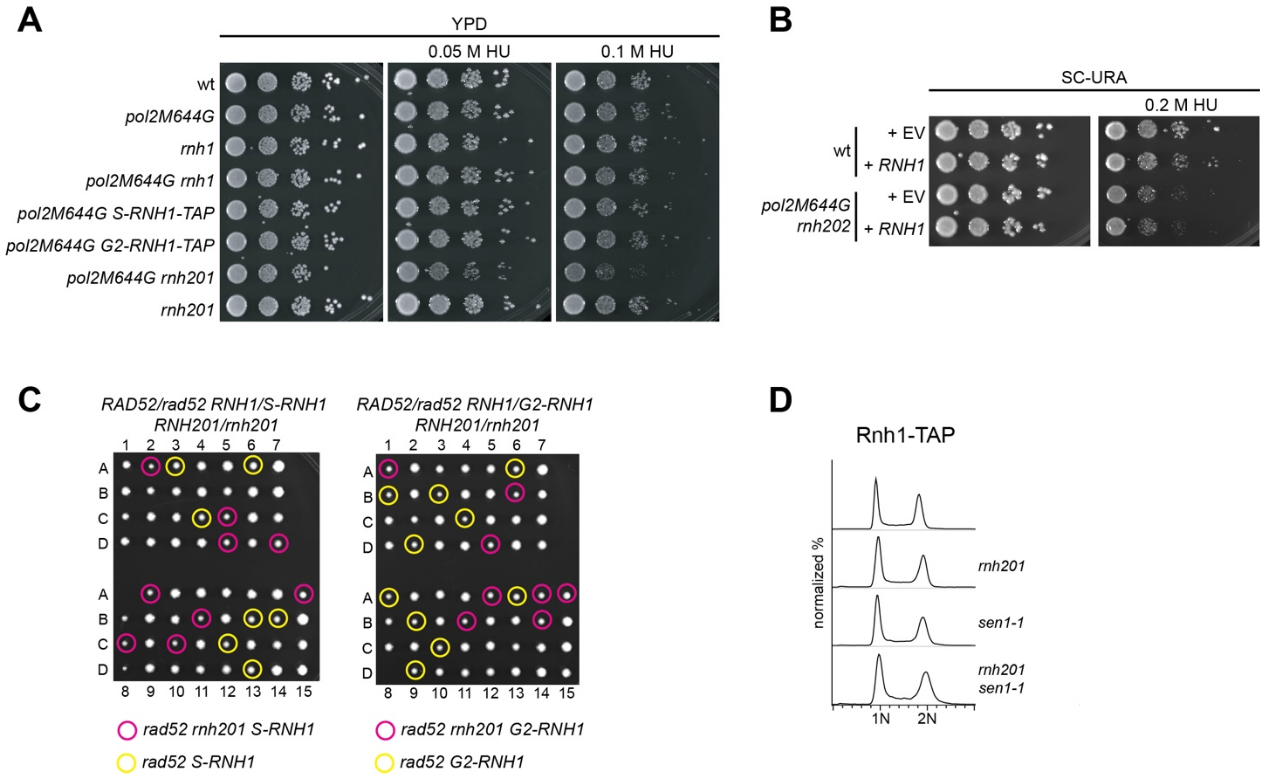


**Figure S3. RNase H1 is specific for R-loop removal. Related to Figure 5.**

(A) A tenfold serial dilution of the indicated strains was spotted onto rich medium with or without 0.05 or 0.1 M HU. Images were taken after 2 days growth at 30˚C.

(B) A tenfold serial dilution of the indicated strains transformed with the indicated plasmids was spotted onto SC-URA plates with or without 0.2 M HU. Images were taken after 3 days growth at 30˚C. EV = empty vector.

(C) Heterozygous diploids with the cell cycle alleles of *RNH1* were dissected in in the presence and absence of both *RAD52* and *RNH201.*

(D) Flow cytometry analysis of DNA content of samples shown in Figure 5E.
